# Susceptibility-matched padding improves the quality of cervical and lumbar spinal fMRI

**DOI:** 10.64898/2025.12.05.692545

**Authors:** Olivia S. Kowalczyk, Matthew A. Howard, David J. Lythgoe

## Abstract

Spinal cord functional magnetic resonance imaging (fMRI) has advanced significantly in recent years, revealing insights into the function of somatosensory and motor systems. However, the complex environment of the spinal cord induces unique sources of noise, limiting the quality of spinal fMRI recordings. Various hardware and software solutions have been proposed to address these challenges. Among them, susceptibility-matched padding has gained popularity due its low cost, ease of use, and effectiveness in reducing static B_0_ field inhomogeneities, which are a major source of artefacts in spinal fMRI. Despite anecdotal evidence, the impact of susceptibility-matched padding on the quality of spinal cord fMRI has not been assessed systematically. We investigated the effects of non-protonated perfluorocarbon liquid-filled padding (SatPad) on B_0_ field homogeneity and functional echo-planar imaging (EPI) in cervical and lumbar spinal cord in 10 healthy volunteers. Participants underwent two resting-state fMRI scanning sessions, one per cord section. Within each session they were scanned with and without SatPad in a pseudo-randomised order. The use of SatPad increased B_0_ field homogeneity and improved functional image quality metrics, including temporal signal-to-noise ratio, timeseries coefficient of variation, DVARS, and ghosting artefacts. While both cervical and lumbar cord data benefited from the use of SatPad, greater effects were observed in the cervical cord. These findings provide a compelling basis for integrating susceptibility-matched padding into routine spinal fMRI protocols.

## 1 Introduction

The field of spinal functional magnetic resonance imaging (fMRI) has developed rapidly over the last two decades, offering substantial insights into the function of somatosensory and motor systems [1,2]. Nevertheless, despite technological and analytical progress, acquiring high quality fMRI data from the spinal cord remains challenging [1].

Several intrinsic characteristics of the spinal cord contribute to the difficulties in recording fMRI signals. The spinal cord is a relatively small and deep-seated structure, surrounded by many different tissue types. This means not only that many currently available MRI surface coils can have lower sensitivity to spinal cord signals but also that these signals can be compromised by differing magnetic susceptibility profiles of surrounding tissues [1–3]. The spinal cord is also in close proximity to many physiologically active regions, such as cardiac, respiratory, and digestive systems, which can further contribute to signal distortions [4,5]. In recent years, the field has seen advancements in purpose-made radiofrequency coils, optimised acquisition protocols, and advanced shimming procedures, all resulting in improved quality of spinal fMRI data [6–8]. Despite this progress, the quality of spinal fMRI recordings is not always consistent and lags behind that of brain fMRI.

Another solution to the challenges of spinal fMRI is the use of commercially available non-protonated perfluorocarbon liquid-filled pads (e.g. SatPad, https://satpads.com). SatPad technology has been suggested to improve the quality of MRI data primarily by reducing static magnetic field (B_0_) inhomogeneities and improving fat suppression [9,10], with similar observations reported using other susceptibility-matched materials [11,12]. Additionally, the padding offers the potential to reduce motion artefacts through improved in-scan participant comfort, gentle soft tissue compression, and optimal body positioning.

Given that areas such as the neck (cervical cord) and torso (thoracic/lumbar cord) are particularly prone to susceptibility artefacts due to their irregular contours, non-uniform soft tissue distributions, in addition to distortions induced by proximity to other physiologically active systems, cervical and lumbar spinal fMRI, in particular, may benefit from the use of susceptibility-matched padding. Although the use of SatPad and similar solutions has been widely adopted in the spinal fMRI community [1,2], apart from two conference abstracts showing promising results [13,14], their efficacy in improving the quality of spinal fMRI signal has not been systematically assessed.

Here we tested the effects of SatPad on two regions of the spinal cord, cervical and lumbar, in a group of 10 healthy volunteers using an echo-planar imaging (EPI) sequence with custom static second order and dynamic slice-specific xyz-shimming [8]. Volunteers’ cervical and lumbar cords (in separate sessions) were scanned with and without SatPad in a pseudo-randomised order. We predicted the B_0_ field would be more homogenous and the quality of functional data would be improved with SatPad in place. Specifically, we hypothesised that SatPad will lead to higher temporal signal-to-noise ratio (tSNR), lower signal variability, less in-scan motion, and fewer ghosting artefacts.

## 2 Material and methods

### 2.1 Participants

Ten healthy volunteers (6 females, 4 males) aged between 24 and 34 years (mean ± standard deviation = 29 ± 4 years) participated in this study. Participants had no history of spinal abnormalities, spinal trauma, nor any MRI contraindications. Ethical approval was acquired from the local research ethics committee (approval number HR-20/21– 21,138) and written informed consent was given by all volunteers.

### 2.2 Procedure

Volunteers underwent two scanning sessions: one focussed on the cervical spinal cord and one on the lumbar spinal cord. Within each session, participants were scanned with (SatPad+) and without SatPad (SatPad-) in a pseudo-randomised order. Each set of scans (SatPad+ and SatPad-) included functional and structural data acquisitions. Cervical spinal scans involved the use of the Neck & Brachial Plexus SatPad set, where the neck pad was fitted as a collar around the volunteer’s neck while the brachial plexus pad was rested on the volunteer’s brachial plexus and fixed with velcro straps to the neck pad. Lumbar spinal cord scans used the Back & Abdomen SatPad, positioned on top of the volunteer’s abdomen.

### 2.3 MRI acquisition

Data were acquired at the Centre for Neuroimaging Sciences, King’s College London using a 3T GE SIGNA Premier XT System (General Electric, Chicago, Illinois) equipped with a 48-channel head and neck coil, a 60-channel posterior array coil, and a 30-channel AIR anterior array coil. T2-weighted structural data was acquired using a sagittal 3D CUBE sequence over 56 slices (repetition time (TR) = 2.5 s, echo time (TE) = 124.3 ms, echo train length = 78, flip angle = 90°, field of view (FOV) = 300 mm, acquisition matrix = 320 × 320, slice thickness = 0.8 mm). This acquisition was based on Cohen-Adad et al. [15] with the FOV increased to 300 mm. For cervical scans, the acquisition covered the whole brain and cervical cord, with the inferior-most slice positioned midway through the T2 vertebra. For lumbar scans, the inferior-most slice was anchored at the bottom of the L5 vertebra covering inferior sections of thoracic cord and the whole lumbo-sacral cord.

Static first- and second-order shims were optimised prior to resting-state scanning. A spectral-spatial excitation pulse was used to excite only tissue water and slice-specific linear shims were implemented by adding 0.6 ms duration x, y, and z-gradient lobes after the excitation pulse. Second-order static and dynamic xyz-shimming were optimised over elliptical regions of interest (ROIs) covering the cord drawn manually by the researcher present during scanning. The ROIs covered the cord and the surrounding vertebral body for second order shimming, but only cord structures for xyz-shimming. ROIs were saved for later examination of field homogeneity (Section 2.4.2.1). The shimming optimisation was a modification of the procedure described in Tsivaka et al. [8]. Since the GE Premier XT does not have an offset coil (zeroth order shim), the centre frequency was optimised rather than the coil current.

Resting-state functional EPI data was acquired over 100 volumes, preceded by four dummy volumes enabling the signal to reach steady-state. Cervical data comprised 38 sequential descending slices (slice thickness = 4 mm, slice gap = 1 mm), with the inferior-most slices prescribed at vertebral level T2 and covering the whole cervical spinal cord and the brainstem (TR = 2.5 s, TE = 30 ms, flip angle = 90°, ASSET factor = 2, FOV = 180 mm, acquisition matrix = 96 × 96, reconstruction matrix = 128 × 128, in-plane reconstructed voxel size = 1.41 mm × 1.41 mm). Lumbar data was acquired over 23 sequential slices in descending order (slice thickness = 4 mm, slice gap = 1 mm), with the inferior-most slices prescribed at the base of vertebral level L1 and covering the whole lumbar spinal cord (TR = 1.6 s, TE = 30 ms, flip angle = 90°, ASSET factor = 2, FOV = 150 mm, acquisition matrix = 96 × 96, reconstruction matrix = 128 × 128, in-plane reconstructed voxel size = 1.17 × 1.17 mm). A multi-slice field map was also acquired for the assessment of field homogeneity, with the same slice prescription as the subsequent resting-state image and a frequency range of ±1000 Hz.

### 2.4 Data analysis

The code used to conduct the analyses described here can be accessed at https://github.com/oliviakowalczyk/satpad.

#### 2.4.1 Preprocessing

Minimal preprocessing was performed on the data to avoid artificially inflating outcome measures. Data were processed using Spinal Cord Toolbox (SCT) version 6.5 [16], AFNI’s *3dWarpDrive* [17,18], and FSL version 6.0.7.4 [19,20]. Visual quality assurance was performed on raw data and at each stage of processing.

For cervical cord data, preprocessing began with the removal of brainstem structures at the level of the odontoid process using *fslroi*. All subsequent preprocessing steps were the same for cervical and lumbar data, unless specified otherwise, and performed in parallel.

Functional data were motion-corrected for slice-wise x- and y-translations using an in-house implementation of AFNI’s *3dWarpDrive* following the steps in the Neptune Toolbox (https://neptunetoolbox.com/). Nearest neighbour interpolation was used to avoid modifying the noise characteristics of the data and artificially inflating our measures.

Warping parameters for spatial normalisation were determined by segmenting and registering the functional data to the PAM50 spinal cord template [21], via an intermediary subject-specific T2-weighted 3D volume. Specifically, *sct_deepseg* [22] was used to segment the cord from the cerebrospinal fluid on the temporal mean of motion-corrected functional data and on the T2-weighted structural image. Where necessary for accurate functional data segmentation, manual adjustment of cord masks was performed in FSLeyes [23]. Warping parameters for registration of functional data to the PAM50 template were created by combining warp parameters from: (1) registering the structural T2-weighted image to the functional data and (2) registering the subject-specific T2-weighted image to the PAM50 T2-weigthed template via *sct_register_to_template* [21]. Intervertebral discs were labelled manually and used for cervical cord registration, along with cord masks. Disc-based registration, however, is not suitable for lumbar cord registration, given the smaller size of the cord and greater variability in vertebral body-to-spinal level alignment in that region [24]. Consequently, lumbar cord registration used manual labels placed at the conus medullaris and intervertebral disc T9-T10, along with cord masks (https://spinalcordtoolbox.com/stable/user_section/tutorials/registration-to-template/lumbar-registration.html). Warps were applied to tSNR maps via *sct_register_multimodal* [21] using nearest neighbour interpolation to avoid smoothing the data. tSNR maps in PAM50 space were used for illustration purposes only.

#### 2.4.2 Image quality metrics

The image quality metrics described below were derived from motion-corrected functional data, with the exception of field homogeneity, which used the field map. Metrics were calculated in native space separately for the cervical and lumbar cord.

##### 2.4.2.1 Field homogeneity

The B_0_ field homogeneity was assessed using the standard deviation of the spherical harmonic fit produced during optimisation of the static shims. The standard deviation (in Hz) was calculated across all voxels within the ROI used for shim optimisation (including those with a value of zero).

##### 2.4.2.2 tSNR

tSNR describes the fluctuations in signal-to-noise profile of data over time and is often used in fMRI to describe the stability of blood oxygen level-dependent signal over the course of a functional scan. tSNR maps were created on motion-corrected data by dividing the temporal mean of the functional image by its standard deviation. Mean tSNR was extracted for the whole cervical (including the first thoracic segmental level; C1-T1) and lumbar (L1-L5) cords, as well as for individual spinal segmental levels in each cord section (C1-C8 and L1-L5). Subject-specific cord masks generated during preprocessing (Section 2.4.1) were used to extract whole-cord mean tSNR, while relevant spinal segmental level masks from the PAM50 atlas warped into native space were used to extract average tSNR within each spinal segment (C1-C8 and L1-L5).

Groupwise tSNR maps were created in PAM50 space by averaging subject-specific maps and masking the resultant image with a whole-cord PAM50 mask. This groupwise map was used for visualisation only.

##### 2.4.2.3 Signal variability

To assess the overall signal variability throughout the functional acquisition, the coefficient of variation of the timeseries was calculated. Coefficient of variation was estimated from motion-corrected data by dividing the standard deviation of the functional image by its mean. Mean coefficients of variation were extracted for the whole cervical and lumbar cords.

##### 2.4.2.4 DVARS

The root mean square of the temporal derivative of the data (DVARS) was used to quantify the rate of framewise signal change. DVARS was calculated using ComputeDVARS from version 1.1.0 of the *nipype*.*algorithms*.*confounds* module [25] within the subject-specific cord masks generated during preprocessing (Section 2.4.1). Mean DVARS was computed for cervical and lumbar cord separately.

##### 2.4.2.5 Motion

The effects of participant motion were characterised using framewise displacement. Given that cord translations in the rostro-caudal plane and rotations around all planes are negligible [5,26], an adapted version of framewise displacement calculation was employed incorporating only x- and y-translations obtained during motion correction (Section 2.4.1): *FD*_*i*_ = |Δ*d*_*ix*_| + |Δ*d*_*iy*_|, where Δ*d*_*ix*_ = *d*_(*i*−1)*x*_ − *d*_*ix*_ and Δ*d*_*iy*_ = *d*_(*i*−1)*y*_ − *d*_*iy*_. Mean framewise displacement was calculated for cervical and lumbar cord separately.

##### 2.4.2.6 Ghosting artefacts

The presence of image ghosting, a common artefact in spinal fMRI acquisitions, in functional data was assessed qualitatively. The number of slices showing ghosting artefacts was counted for each participant’s cervical and lumbar functional acquisitions.

#### 2.4.3 Statistical analysis

The effects of SatPad on image quality metrics, including field homogeneity, whole-cord and segmental tSNR, timeseries coefficient of variation, DVARS, framewise displacement, and ghosting were assessed using paired-samples t-tests conducted in Python version 3.12.9 with the Pingouin package version 0.5.5 [27]. All comparisons were performed for the cervical and lumbar cord separately.

## 3 Results

### 3.1 Field homogeneity

SatPad+ led to a 29% decrease in the standard deviation of the magnetic field within the ROI used to optimise the second-order static shim for the cervical cord (t(9) = 3.60, *p* = 0.006, 95% CI [7.93, 34.72], Hedges *g* = 1.20). The standard deviation of the field in the lumbar cord was 11% lower for SatPad+, but this was not statistically significant (t(9) = 2.04, *p* = 0.073, 95% CI [−0.62, 11.69], Hedges *g* = 0.34; Figure 1A).

**Figure 1.**
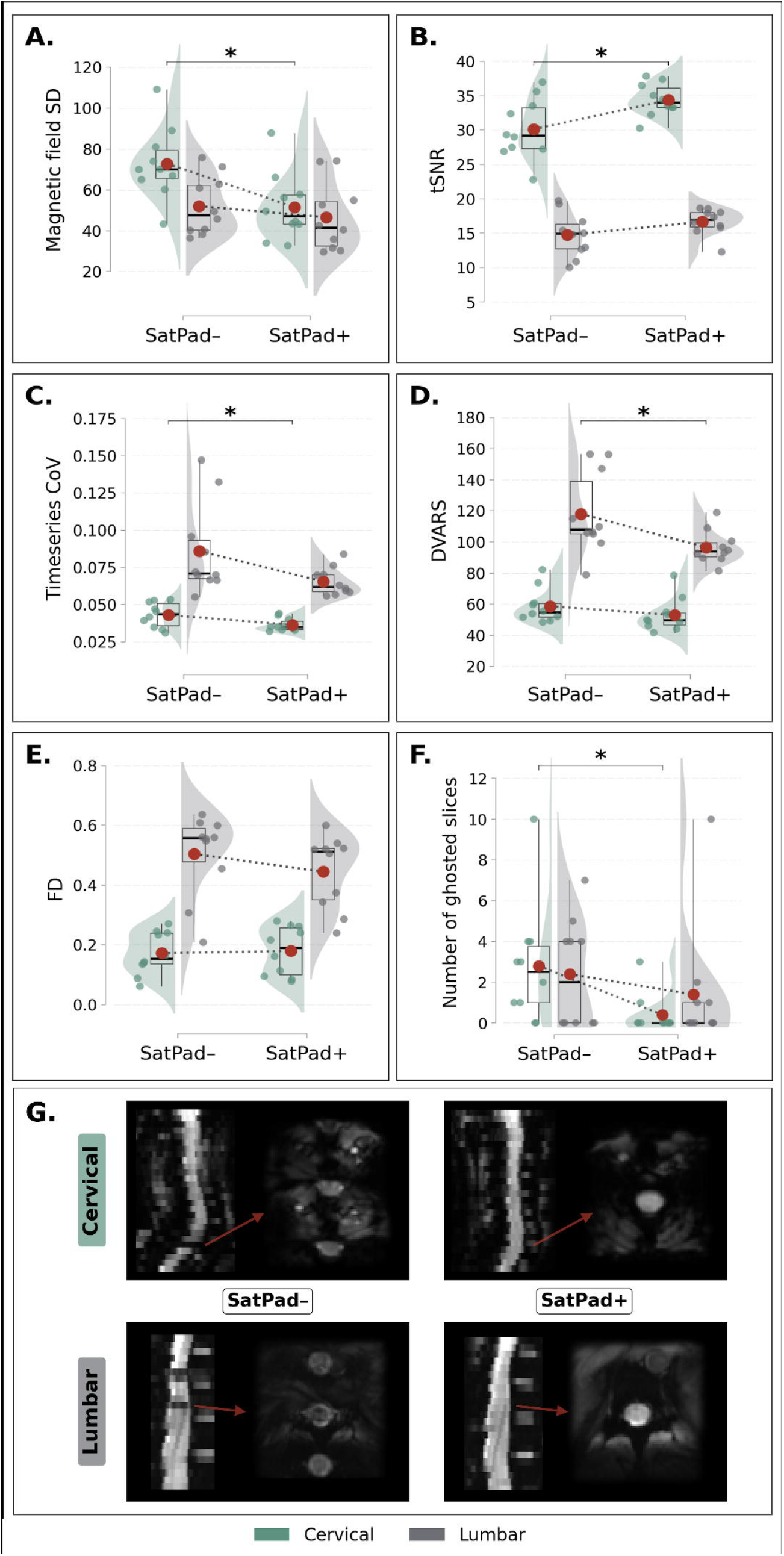
Comparisons of SatPad- and SatPad+ on measures estimated on the whole section of cervical (teal) and lumbar (grey) cord. Boxplots mark the median (thick black line), interquartile range (box), and minimum and maximum values (whiskers). Red dots mark the mean, with dashed lines indicating the direction of the effect. **A**. Magnetic field homogeneity during static shim optimisation (expressed as standard deviation). **B**. Temporal signal-to-noise ratio. **C**. Timeseries coefficient of variation. **D**. DVARS. **E**. Framewise displacement (in millimetres). **F**. Number of slices showing ghosting artefacts. **G**. Exemplary EPI images in native space showing the effects of SatPad on ghosting artefacts in cervical (top row) and lumbar (bottom row). CoV, coefficient of variation, DVARS, difference of variance; FD, framewise displacement; SatPad-, acquired without SatPad; SatPad+, acquired with SatPad; SD, standard deviation; tSNR, temporal signal-to-noise ratio. \**p* < 0.05

### 3.2 tSNR

SatPad+ was associated with a 14% increase in tSNR extracted from the whole cervical spinal cord (*t*(9) = 2.78, *p* = 0.021, 95% CI [−7.72, −0.79], Hedges *g* = 1.15). A similar 13% increase in whole-cord tSNR was also observed in the lumbar cord, but it was not statistically significant (*t*(9) = 1.54, *p* = 0.159, 95% CI [−4.77, 0.91], Hedges *g* = 0.69; Figure 1B). When assessing individual spinal cord segments, statistically significant gains in tSNR were observed at spinal segmental level C3 (24% increase, *t*(9) = 2.38, *p* = 0.042, 95% CI [−14.35, −0.35], Hedges *g* = 0.98), C4 (31% increase, *t*(9) = 2.77, *p* = 0.022, 95% CI [−15.26, −1.53], Hedges *g* = 1.24), and C7 (17% increase, *t*(9) = 5.34, *p* < 0.001, 95% CI [−7.56, −3.09], Hedges *g* = 1.02). While SatPad+ use was associated with higher tSNR across remaining cervical segmental levels and all lumbar segmental levels, these differences did not reach statistical significance (all *p* < 0.05; Figure 2).

**Figure 2.**
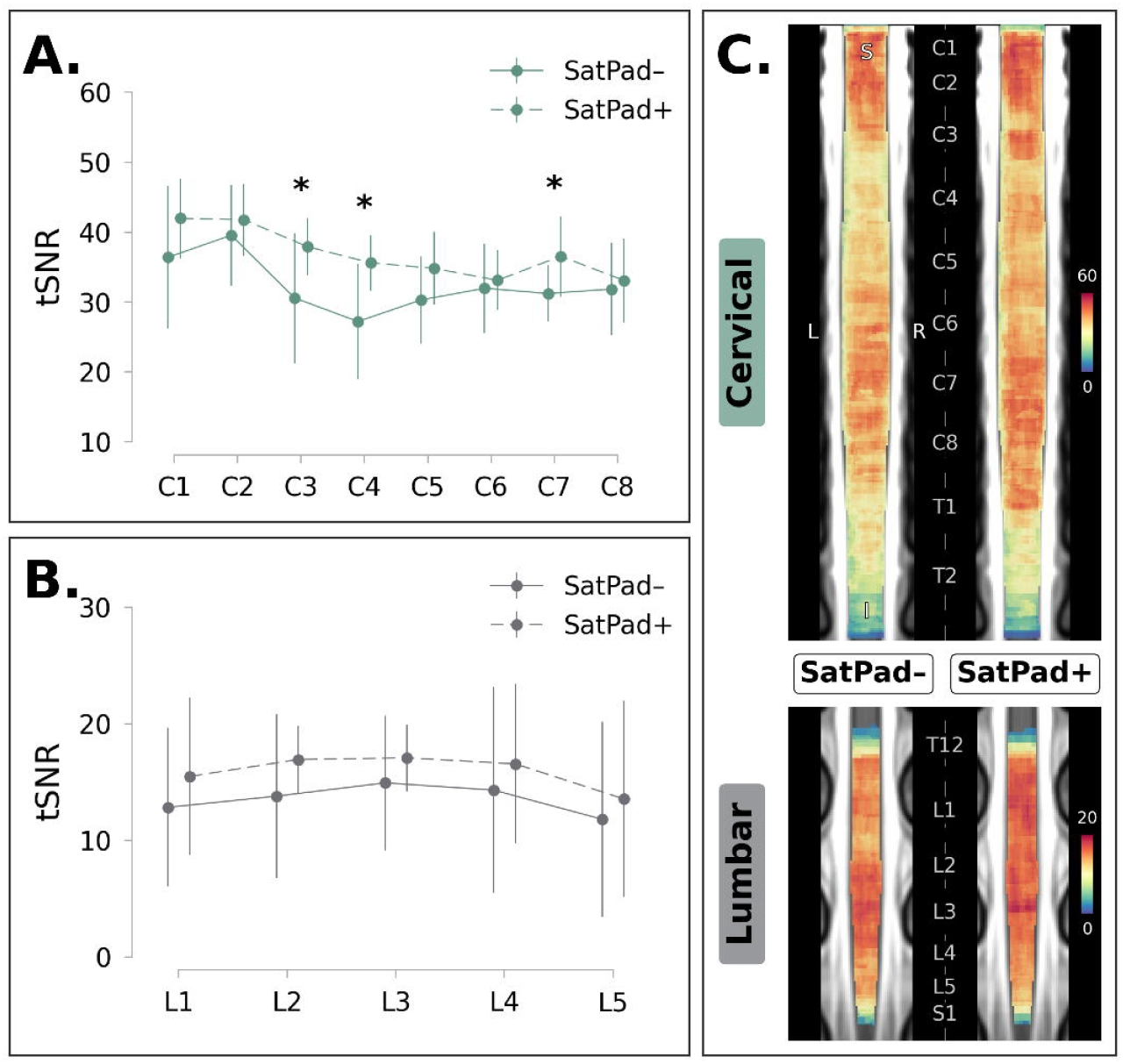
Temporal signal-to-noise ratio (tSNR) comparisons between SatPad- and SatPad+ in distinct regions of the cervical (teal) and lumbar (grey) cord. **A**. Mean (± SD) tSNR in each cervical spinal cord segmental level. **B**. Mean (± SD) tSNR in each lumbar spinal cord segmental level. **C**. Group-level maps showing tSNR values in cervical (top panel) and lumbar (bottom panel) in PAM50 space. Labels denote spinal segmental levels. tSNR, temporal signal-to-noise-ratio; SatPad-, acquired without SatPad; SatPad+, acquired with SatPad. SD, standard deviation. \**p* < 0.05

### 3.3 Signal variability

SatPad+ was related to a 15% reduction in timeseries coefficient of variation in the cervical spinal cord (*t*(9) = 2.34, *p* = 0.044, 95% CI [<0.01, 0.01], Hedges *g* = 0.95) and 24% reduction in lumbar spinal cord, albeit not statistically significant (*t*(9) = 1.86, *p* = 0.097, 95% CI [0.00, 0.05], Hedges *g* = 0.87; Figure 1C).

### 3.4 DVARS

DVARS was decreased by 10% in the SatPad+ condition in cervical cord, although this did not reach statistical significance (t(9) = 2.22, *p* = 0.054, 95% CI [−0.12, 1.55], Hedges *g* = 0.50), and by 18% in the lumbar cord (t(9) = 2.32, *p* = 0.046, 95% CI [0.53, 42.35], Hedges *g* = 1.03; Figure 1D).

### 3.5 Motion

Although SatPad+ was related to small changes in framewise displacement across both cervical (4% increase, t(9) = 0.42, *p* = 0.686, 95% CI [−0.05, 0.03], Hedges *g* = 0.09) and lumbar cord (12% decrease, t(9) = 1.74, *p* = 0.116, 95% CI [−0.02, 0.14], Hedges *g* = 0.43), these effects were not statistically significant (Figure 1E).

### 3.6 Ghosting artefacts

Ghosting artefacts were reduced by 86% in the SatPad+ condition in the cervical cord (t(9) = 3.50, *p* = 0.007, 95% CI [0.85, 3.95], Hedges *g* = 1.05), and by 42% in the lumbar cord, although lumbar effects did not reach statistical significance (t(9) = 0.90, *p* = 0.393, 95% CI [−1.52, 3.52], Hedges *g* = 0.33; Figure 1F and Figure 1G).

## 4 Discussion

This study systematically characterised the effects of non-protonated perfluorocarbon liquid-filled pads (SatPad) on the quality of cervical and lumbar spinal cord fMRI. SatPad use increased B_0_ field homogeneity and was associated with meaningful improvements in functional image quality, as indexed by tSNR, timeseries coefficient of variation, DVARS, and ghosting artefacts. Although both cervical and lumbar cord data benefited from the use of SatPad, gains were more pronounced in the cervical cord.

Field inhomogeneity is a key driver of many artefacts and distortions limiting the quality of spinal fMRI. Consistent with previous reports [9–11,14], we observed that the use of susceptibility-matched padding substantially improved the homogeneity of the B_0_ field, with greater effects in the cervical cord. SatPad, and other analogous devices, have similar magnetic susceptibility to biological tissues, and thus displace susceptibility-driven field gradients at air-tissue boundaries away from the imaged structure [12]. The larger improvements in field homogeneity in cervical, compared to lumbar cord, thus likely reflect the closer proximity of air-tissue boundaries in the neck compared to the abdomen, where somewhat more homogenous soft tissues surround the cord. Several other anatomical factors may further contribute to lower SatPad efficacy in the lumbar cord, including more variable sources of physiological noise, larger vertebral bony segments, and greater distance between the padding counteracting those effects and the cord [28]. Additionally, cervical, but not lumbar SatPad, can improve in-scanner body positioning and reduce spinal cord curvature. This may offer additional improvements to field homogeneity by enabling better shim volume alignment with the cord and minimising the inclusion of air-tissue boundaries [15]. Such effects may be less pronounced in ROI-based shimming approaches, such as the one used here, where air-tissue boundaries are outside of the ROIs.

The biggest impacts of SatPad on functional data quality were in reducing ghosting artefacts and increasing tSNR. Ghosting is common in spinal cord fMRI, particularly in EPI acquisitions which have become widely adopted by the field due to their fast sampling capacity and high signal sensitivity [1,2,29]. Ghosting artefacts arise from B_0_ field inhomogeneities and can be further exacerbated by time-varying motion. This is especially relevant to the spinal cord, where periodic movement is induced by surrounding physiological processes, including respiration, heartbeat, cerebrospinal fluid pulsation, swallowing, and digestion [28]. These compound effects of noise are therefore more pronounced in cord regions proximal to physiologically active organs, such as lower cervical, thoracic, and lumbar segments [30,31]. While our data show the largest tSNR boost with SatPad in rostral regions, improvements were also observed in more caudal cervical segments, paralleling previous reports [13], as well as in the lumbar cord. Given the technical challenges of imaging these segments, susceptibility-matched padding may be particularly advantageous in such cases.

SatPad also led to decreases in timeseries coefficient of variation and DVARS, indicating improved signal stability across lumbar and cervical cord. Notably, framewise displacement showed no reduction in cervical cord and only marginal decreases in lumbar cord, suggesting that the functional signal quality improvements observed with SatPad are unlikely to be explained by movement suppression. Nonetheless, our sample was small and comprised experienced MRI participants; studies recruiting scanner-naïve individuals may still benefit from the added stabilisation offered by SatPad. The observed improvements in functional signal quality and stability in cervical and lumbar cord align with expectations that susceptibility-matched padding would lessen the impact of some of the specific challenges related to spinal imaging. These effects are likely driven by increases in B_0_ field homogeneity as demonstrated here and in previous studies [9,10,14]. Beyond improving field homogeneity, SatPad can facilitate optimal body positioning with minimal spine curvature, particularly for cervical cord imaging, enabling perpendicular axial slice prescriptions that minimise partial volume effects and ensuring robust performance of image registration algorithms [1]. By mitigating many of the factors leading to artefacts and signal instability, susceptibility-matched padding should improve the reliability of spinal fMRI [32,33], and support longitudinal and multi-site investigations.

Although we focussed on spinal cord functional EPI at 3 Tesla, our findings may extend to other field strengths and other sequences prone to B_0_ inhomogeneity artefacts, such as gradient echo and accelerated acquisitions [34]. Supporting this potential generalisability, susceptibility-matched padding has shown benefits in structural and diffusion-weighted imaging of the head and neck [35,36]. Improved field homogeneity has also been observed in cervical spinal imaging at 7 Tesla [10], although detailed characterisation of the effects on functional image quality is lacking.

## 5 Conclusions

Our findings demonstrate the utility of SatPad in reducing B_0_ field inhomogeneities and enhancing spinal fMRI data quality across several metrics. These improvements hold potential for more reliable interpretation of spinal fMRI data, facilitating its broader application in both basic and clinical research.

## Glossary

B_0_: static magnetic field
CI: confidence interval
CoV: coefficient of variation
DVARS: root mean square of the temporal derivative of the data
EPI: echo-planar imaging
FD: framewise displacement
fMRI: functional magnetic resonance imaging
FOV: field of view
MRI: magnetic resonance imaging
PAM50: Polytechnique Montreal, Aix-Marseille Université and Montreal Neurological Institute
ROI: region of interest
SatPad-: data acquired without SatPad
SatPad+: data acquired with SatPad
SCT: Spinal Cord Toolbox
SD: standard deviation
TE: echo time
TR: repetition time
tSNR: temporal signal-to-noise ratio

## Acknowledgements

We thank all volunteers for their participation in the study.

